# In search of a brain microbiome: A machine learning search pipeline for electron microscopy images of brain tissue

**DOI:** 10.1101/2022.07.12.499807

**Authors:** Jordan K. Matelsky, Celina Shih, Khalil Hijazi, Erik C. Johnson

**Affiliations:** Johns Hopkins University Applied Physics Laboratory, Laurel, Maryland, United States; University of Pennsylvania Department of Bioengineering, Philadelphia, Pennsylvania, United States

**Author notes:** Research was sponsored by the Army Research Office and was accomplished under Grant Number W911NF-20-1-0166. The views and conclusions contained in this document are those of the authors and should not be interpreted as representing the official policies, either expressed or implied, of the Army Research Office or the U.S. Government. The U.S. Government is authorized to reproduce and distribute reprints for Government purposes notwithstanding any copyright notation herein. The team would like to thank Nathan Boggs, Monique Beaudoin, Matthew Hart, and Brock Wester for their valuable guidance. Finally we would like to thank the researchers who made their data public and enabled this work.

## Abstract

The healthy human brain has long been considered a sterile environment, with the blood brain barrier preventing the formation of a bacterial brain microbiome. Recent electron microscopy (EM) imaging of brain tissue has, however, provided the first preliminary evidence of bacteria in otherwise healthy brain slices. Whether due to contamination, disease, or a previously unknown relationship of bacteria to healthy brain tissue, novel tools are needed to detect and search for bacteria in nanoscale, volumetric EM images. While computer vision tools are widely used in cell segmentation and object detection problems in EM imaging, no bacteria detection tool or dataset exists. Overcoming the rarity of training data, this work presents the first pipeline for training a bacteria detection network for EM images, leveraging existing deep networks for object detection. A deployment and proofreading pipeline is presented, along with characterization of deployment to public EM image datasets. While bacteria in healthy brain tissue were not discovered in this work, this tool presents an opportunity for large scale bacteria search in EM imaging for both scientific discovery and experimental quality control, and serves more generally as a framework for sparse object detection in large imagery datasets.

## I. INTRODUCTION

While the vertebrate brain has long been thought of as a sterile environment, preliminary manual review of electron microscopy (EM) imagery of mammalian cortex has suggested the presence of bacteria in both healthy as well as diseased brain tissue. Early results from germ-free controls suggest that this was not simply due to sample contamination [1]. This finding conflicts with the widely-held neuroscientific understanding that the blood-brain barrier, a cellular boundary which isolates the central nervous system, is impermeable to bacteria in a healthy individual [2], [3], [4]. If these early findings are confirmed, they may radically alter our understanding of disease in the brain. The major barriers preventing large-scale analysis are (1) the immense scale of nanoscale EM imagery (a single cubic millimeter of tissue can reach petabyte scale [5], [6]); (2) the specialized human expertise required in order to manually scan these datasets for bacteria; and (3) the expense of collecting new datasets.

The scale of the data necessitates an automated segmentation approach: It is not tractable to perform an exhaustive manual review of hundreds of terabytes of raw imagery. Computer vision techniques, including convolutional neural networks, have frequently been used for problems such as cell segmentation or detection of cell substructures in large volumes of bioimagery data [7]. Deep neural networks of different architectures have been extensively used to segment and detect structures such as synapses [8], mitochondria [9], and neurons [10] in EM of different types [11]. These tasks benefit from an abundance of training data — an advantage not shared with the task of bacteria detection in the brain.

We propose a solution to overcome these issues by combining a traditional deep learning object detection network with a training dataset initially seeded by human annotation. We then iteratively refine the output of this algorithm by performing sparse human review and subsequent retraining. This work builds upon existing projects in the storage and processing of large scale EM datasets [12].

Finally, to relieve the requirements to collect new datasets for this research, we relied upon existing public EM datasets, now widely available in online repositories such as BossDB (*bossdb*.*org*). [5] Here, we share our software architecture and the results of our object detection systems.

## II. METHODS

In the presence of sufficient training data, searching for small structures in large EM volumes would be a simple matter of data and infrastructure management. Unfortunately, unlike existing detection networks for synapses and mitochondria, training data for bacterial search is sparse. For this reason, our detection pipeline required a unique approach to generate an adequately sized training dataset.

### A. Training dataset generation

The manually collected dataset consisted of 61 bacterial EM images, curated from various databases, publications, and bacterial image repositories [13], [14], [15], [16], [17], [18], [19], [20], [21]. Because there were no images of bacteria in brain tissue available, we instead searched for endogenous bacteria imagery in other tissue types (such as endothelial or vascular tissue), in order to provide our network with diverse background texture. We captured a wide range of host organisms, tissue types, and bacteria morphologies within the dataset to avoid model biases.

In order to generate training data, we developed a browser-based annotation tool compatible with the Microsoft Common Objects in Context (COCO) object-detection data specification [22] (**Fig. 1**. This tool is designed to read from a user-specified directory of images, and produce JSON annotations compatible with the detection networks used in this work. Our tools support both dense (“mask”) annotation and bounding-box annotation.

**Fig. 1.**
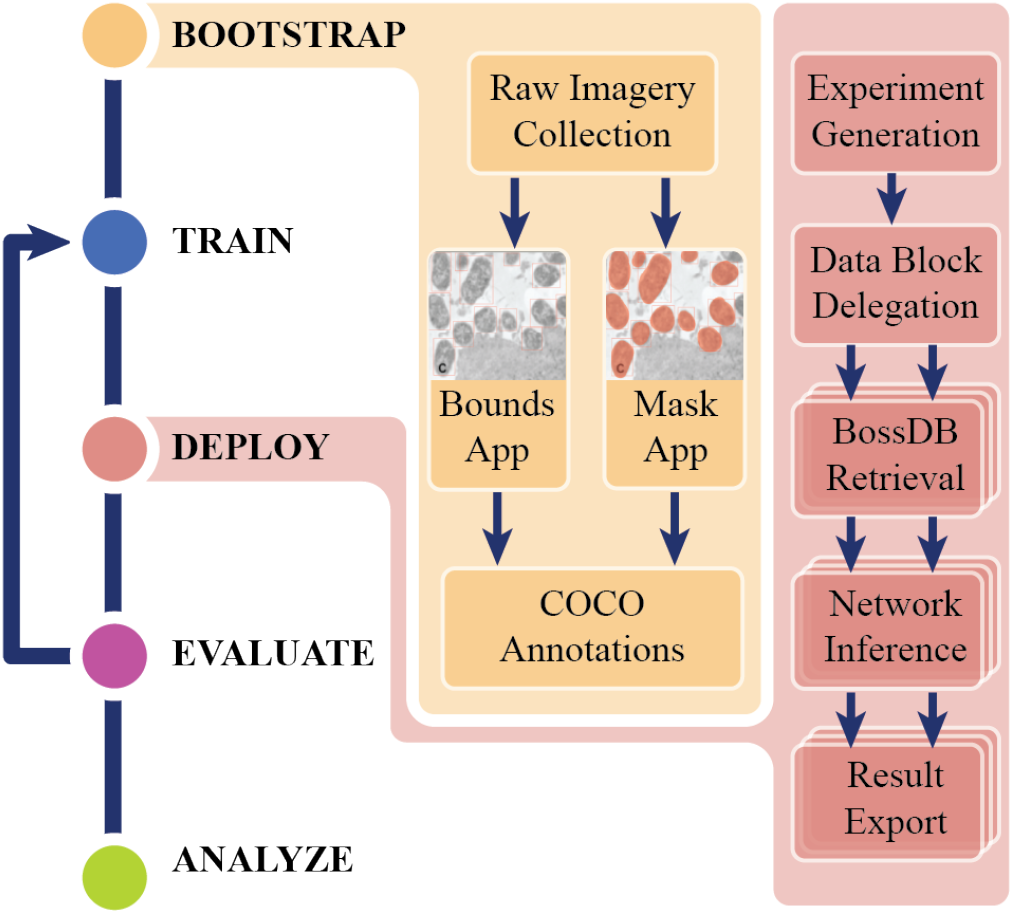
Annotation pipeline for dataset generation. We developed a software suite for dataset generation. Our tools produce COCO-adherent JSON annotations from raw imagery presented in a browser-based user interface. Once a modest training dataset has been generated, the neural network can be trained and its performance may be evaluated. Through expert inspection, errors may be detected and corrected: For example, false-positive inference outputs may be fed back into the training dataset as null examples. This cycle is continued in order to improve performance incrementally, minimizing required human intervention.

### B. Detection Network Training

We built upon the existing *Detectron2* [23], [24] package, which supports a variety of architectures and models for detection and segmentation networks, including both bounding boxes and pixel level segmentation. While not designed specifically for biomedical images, this proven backbone provides extensibility of the proposed approach to new dataset types with minimal engineering effort.

We utilized the *Faster R-CNN* [24] baseline detection network in *Detectron2*. Our manually-annotated bounding boxes served as output labels. (No masks were used during training.) We used a Residual Network backbone with a depth of 50 for all experiments. We computed Mean Average Precision (AP) for all detected objects using ten Intersection over Union thresholds ranging from 0.5 to 0.95, and we used this to determine the best-performing networks on the validation dataset. This was trained with the standard *Detectron2* 3x training schedule [23].

Hyperparameter search was carried out using the small validation set using a manual grid search. The hyperparameters were swept uniformly with *Learning-Rate =* 0.00025 − 0.005 and *Batch-Size =* 1 − 10. The networks with the highest AP on the validation set were selected for deployment. The network used for results below here used a batch size of 1, a base learning rate of 0.00025 and a final epoch count of 600.

### C. Detection Filtering

Due to the density of bacteria in the training dataset and the fact that Fast R-CNN performs multi-scale detections, we expected that these properties would result in a large number of false positives during deployment.

We therefore applied a simple filtering step after detection: The area, in *px*^2^ was computed for each detected object. Objects above a pre-determined area threshold were eliminated, which removed a considerable percentage of false positives (Fig. 4 shows how this threshold can be determined from data in practice).

### D. Deployment Pipeline

We developed an inference pipeline to run many simultaneous deployments using on-premise GPU hardware. We wrote an experiment generator script that output configuration files for image data blocks in JSON format. These configuration files contained the coordinates and specifications for pulling data blocks from BossDB using the *intern* Python library [25]. Our model was then run on each data block to obtain bounding box results (**Fig. 1**). We used several BossDB datasets of various species and brain regions (**Fig. 4**) [26], [27], [28]. We designed this pipeline to be easily adapted to run in a scalable cloud solution for neuroimaging, such as the SABER system [12].

### E. Proofreading and Data Augmentation Pipeline

After deployment over a dataset, results were aggregated and the resulting annotations ordered from highest to lowest confidence. Pre-filtering was used to remove objects over a certain size threshold. These annotations were then fed through a proofreading pipeline, where highest confidence results were presented to a human annotator through the Flask app. The expert then classified the detections as valid or invalid.

We then re-incorporated these annotated frames into new, augmented training datasets, including both negative and positive examples. (It is expected that negative examples are much more prevalent than positive examples in the dataset, so care must be taken not to unbalance the training dataset with negative examples.) This approach enabled us to improve the performance of the network interactively as a dataset is searched. This augmentation loop is shown in **Fig. 1**.

## III. RESULTS

Our best-performing networks were evaluated on the validation dataset. The validation AP was 0.385 (out of a maximum value of 1). This score is likely due to a large number of false positives, particularly related to multi-scale detections. **Fig. 2** shows the resulting training curves and an example result for these data, using a validation image set containing true bacteria in brain EM tissue [1] as well as synthetic imagery.

**Fig. 2.**
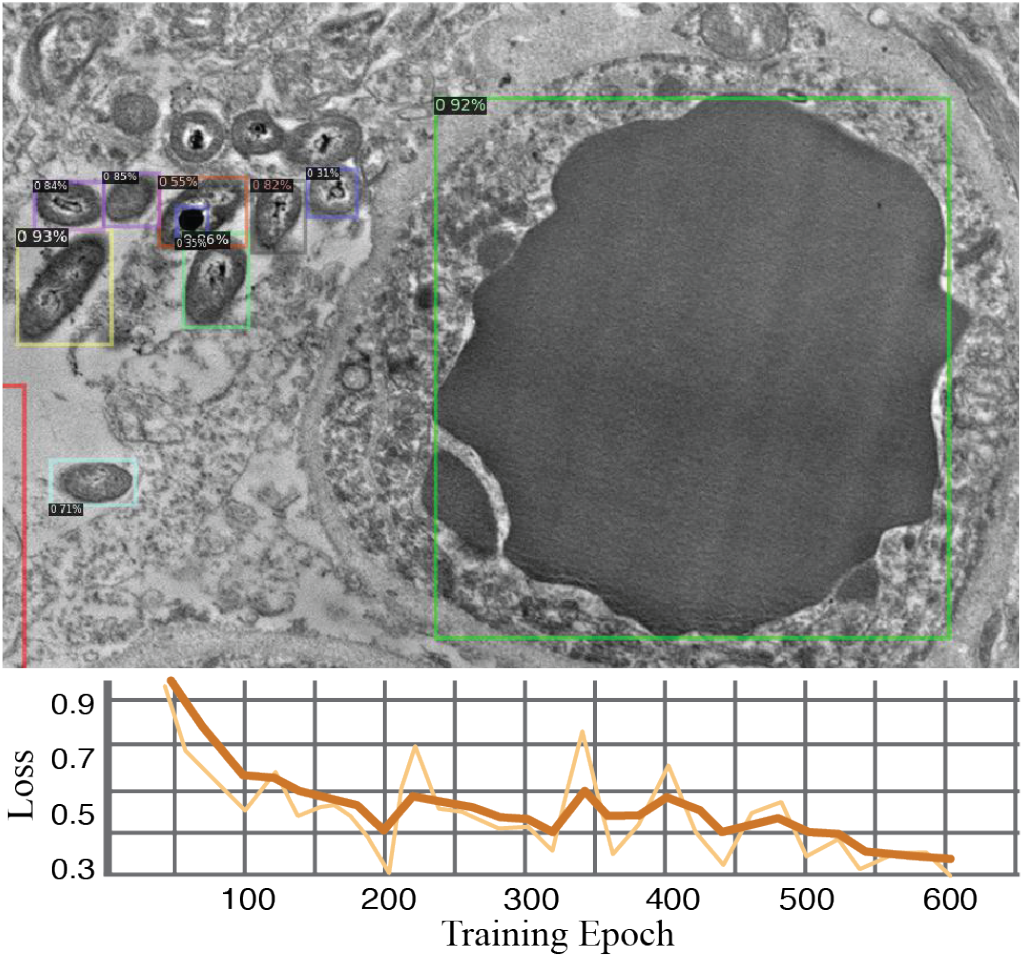
Result of training Fast R-CNN detection network for bacteria detection. Detection on the validation set image containing bacteria in EM images of brain tissue. While there are several large false alarms, these are easily filtered based on size. Below, we show the training loss curve for the network, with the dark line showing running-averaged loss over a sliding window of ten epochs. Above image from Ref. [1].

**Fig. 3.**
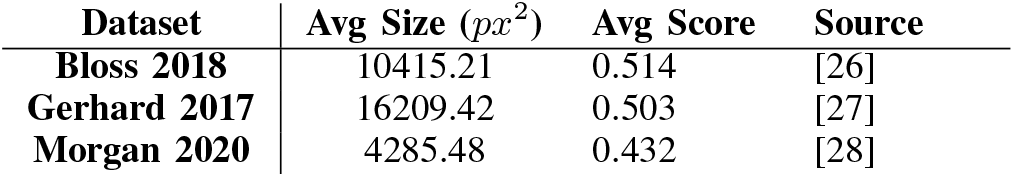
A comparison of dataset results. The average detection size was close to 100, 000*px*^2^, though this varied between datasets. The average score was close to 0.5 for all datasets upon which inference was run.

We ran the highest-performing network described above on several large-scale, 3D EM datasets from BossDB [5], downsampled to 16 × 16 nanometer resolution. Detections tended to fall most frequently in the 10^3^ – 10^4^*px*^2^ scale range (**Fig. 4**). Scores ranged from 0 to 1.0, with the majority of weight close to 0. The average detection size was close to 100, 000*px*^2^, though this varied by dataset (**Table III**). Smaller, false-positive detections tended to be mitochondria, pieces of darkly-staining endoplasmic reticulum, or staining artifacts. Large false-positive detections tended to be blood vessels or slices of red blood cells therein.

**Fig. 4.**
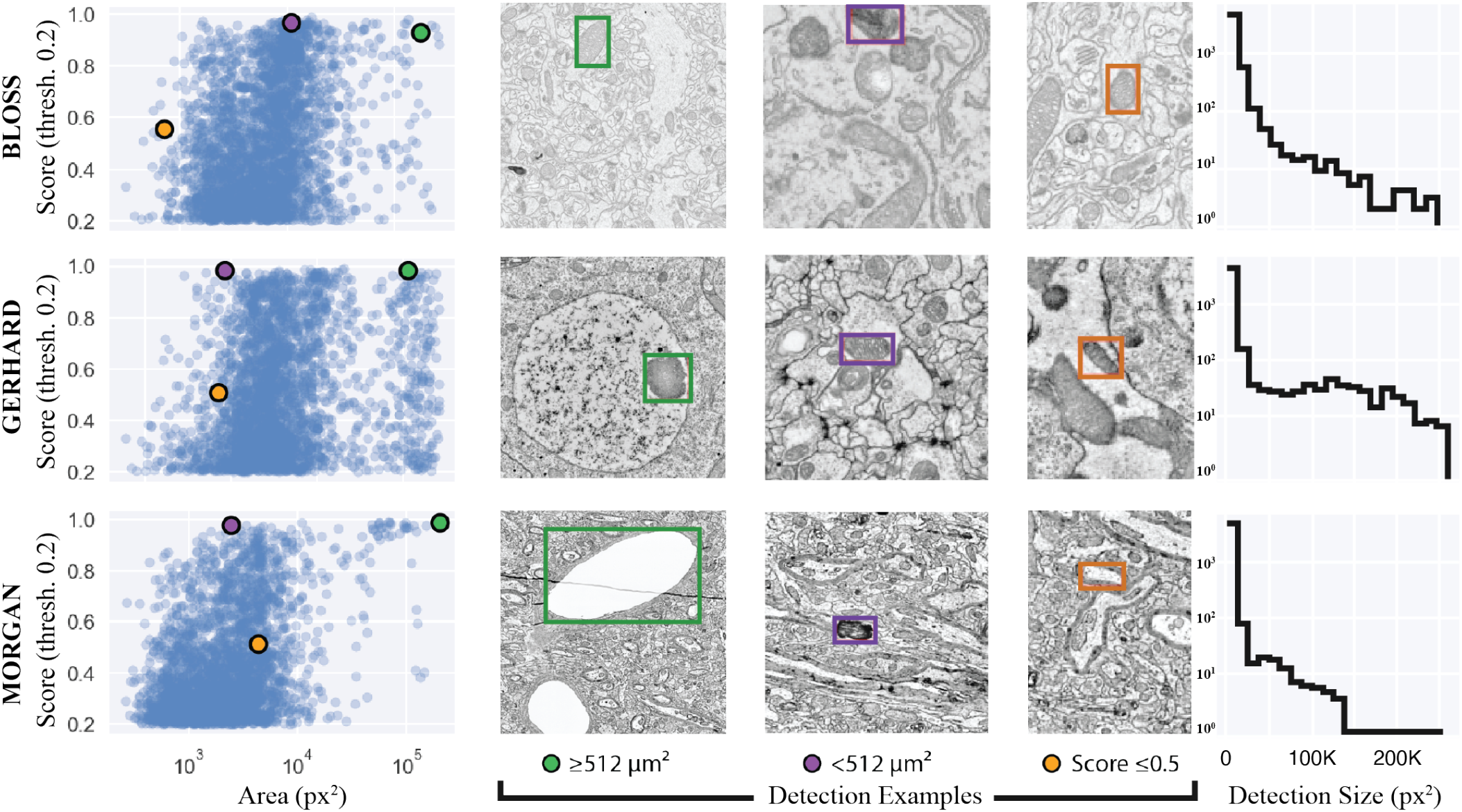
Example detections from public EM datasets. We evaluated our best-performing networks on several public electron microscopy datasets. On the left, we compare the reported confidence scores and detection sizes (in px^2^) of all detections. We show example images from high-confidence large bodies (green), high-confidence small (purple), and low-confidence (orange) detections. On the right, we show detection size histograms for each dataset (*y* axis is on a log scale).

## IV. DISCUSSION

While EM data have very different pixel statistics than natural (photographic) images, and has an inherent 3D structure, we chose to deploy 2D detection networks in order to exploit proven network architectures and simplify the search for small objects in EM into a simple 2D image detection problem. The Faster R-CNN is a proven network architecture capable of detection in a wide range of conditions with a strong invariance to object scale. Though this invariance is a strength in an open-ended search like ours, it can become problematic when taken to an extreme, since bacteria tend to appear at roughly the same scale relative to the imagery (unlike detections in natural images, in which target classes may appear at different distances from the camera). While producing many false positives, this scale invariance property provides robustness to the resolution of EM imaging and size of bacteria types. Future work will consider detection networks specialized for EM applications, especially as improved mask annotations become available.

Our work focused on the technical aspects of dataset construction, network training, and deployment to detect bacteria in EM images. Though there are alternative methods to detect the presence of bacteria in neural tissue, such as genome sequencing, the resolution of EM presents a unique opportunity to confirm the morphology and distribution of bacteria in neuropil. Understanding the locations of bacteria relative to other neural and non-neural structures (such as vasculature) may be critical to understand the possible mechanisms of central nervous system access in different disease or healthy conditions.

We believe that the tools developed here are more generally applicable for any detection task in which few, sparse objects must be found within an expansive image volume, such as in biomedical imagery or satellite imagery. All software and libraries described in this manuscript will be made open-source upon publication.

